# Mice lacking the endocannabinoid-synthesizing enzyme NAPE-PLD exhibit sex-dependent dysregulations in responsiveness to oxycodone and a natural reward

**DOI:** 10.1101/2025.04.03.646908

**Authors:** Taylor J. Woodward, Emily Sizemore, Ananya Balaji, Ada Port, John Hainline, Hasaan Kazi, Serge Luquet, Ken Mackie, Andrea G. Hohmann

## Abstract

The endogenous opioid and endogenous cannabinoid (endocannabinoid) systems are highly interconnected in the context of drug reward. Bioactive lipids known as *N*-acylethanolamines (NAEs), and, specifically, anandamide (AEA), influence several unwanted side effects of opioids, including dependence and tolerance. AEA undergoes degradation by the enzyme fatty-acid amide hydrolase (FAAH), whereas the biosynthesis of AEA *in vivo* is catalyzed by the enzyme *N*-acyl phosphatidylethanolamine phospholipase-D (NAPE-PLD). AEA and FAAH are implicated in opioid reward, but the impact of genetic deletion of NAPE-PLD on responsiveness to opioids remains unknown. Here we explored the role of NAPE-PLD in behavioral sensitivity to the opioid analgesic oxycodone. We evaluated NAPE-PLD knockout (KO) and wild type (WT) mice of both sexes in preclinical assays that assess either opioid-induced psychomotor responses or voluntary oral consumption of oxycodone. In our studies, genetic deletion of NAPE-PLD produced a shift in sexually dimorphic responses to oxycodone. Psychomotor response to oxycodone was reduced in female NAPE-PLD KO mice but not in males. Female NAPE-PLD KO mice consumed more oral oxycodone that female WT mice, while no genotypic differences in consumption were observed in males. Oxycodone consumption also increased the number of striatal ΔFosB positive cells in female WT mice, but not in male WT mice or NAPE-PLD KO mice of either sex. Additionally, NAPE-PLD KO mice of both sexes consumed more sucrose than WT mice. Together, these findings suggest that NAPE-PLD may regulate responses to opioids in a sexually dimorphic manner as the impact of genetic deletion of NAPE-PLD was greater in females than males.

## 1. Introduction

The endogenous opioid and endogenous cannabinoid (i.e., endocannabinoid (eCB)) systems are involved in homeostatic regulation of many physiological processes, including immune function, energy metabolism, and mood. Both systems consist of G-coupled protein receptors (GPCRs) which share many common mechanisms in signal transduction (Gi/o, beta arrestin, MAP kinase, etc.). Both systems are implicated in the reinforcing properties of several drugs of abuse. Activation of opioid and cannabinoid CB_1_ receptors regulate neurotransmitter release via similar mechanisms, and both are densely expressed in brain regions involved in reward processing. Given these shared characteristics, it follows that manipulation of one system could affect the functionality of the other. Preclinical and clinical studies suggest that eCB system-targeting therapies could be potentially useful for treating opioid use disorder (OUD). Despite this promise, limitations in our understanding of how various players in the eCB system affect specific aspects of OUD potentially hinders our ability to advance eCB-based therapeutics.

An interesting and potentially advantageous difference between the opioid system and eCB system lies in the composition and production of their endogenous ligands. While endogenous opioid ligands are peptides, which are created via transcriptional/translational mechanisms, eCB ligands are lipids whose production is regulated by enzymes. The best studied endocannabinoid ligands are 2-arachidonoylglycerol (2-AG) and N-arachidonoylethanolamine (AEA). While the mechanisms surrounding synthesis of 2-AG are well-characterized, the mechanisms of AEA biosynthesis have, until recently, remained incompletely understood. *N*-acylphosphatidylethanolamine-phospholipase D (NAPE-PLD) is the primary enzyme involved in biosynthesis of anandamide and structurally-related *N*-acylethanolamines (NAEs) (Leishman et al., 2016). Despite the current biochemical consensus surrounding NAPE-PLD as the primary enzyme involved in AEA formation, little is currently known about how NAPE-PLD’s activity influences organismal behavior.

Anandamide plays a role in the reinforcing aspects of both opioids and natural rewards. Preclinical studies have largely relied on experimental manipulations that enhance endogenous AEA levels by inhibiting the activity (genetically or pharmacologically) of its degradation enzyme, fatty acid amide hydrolase (FAAH). Negative reinforcement, but not positive reinforcement, from opioid drugs is attenuated by elevating levels of AEA. For example, the FAAH inhibitor URB597 enhanced the extinction of withdrawal-induced conditioned place avoidance, attenuated physical signs of morphine withdrawal, and reversed antinociceptive tolerance to morphine but did not increase reinforcing efficacy of heroin or inhibit the extinction of conditioned place preference (i.e. positive reinforcement), (Fotio et al., 2020; Manwell et al., 2009; Ramesh et al., 2011, 2013; Solinas et al., 2005).

In the present study, we asked whether lowering NAE levels via genetic deletion of NAPE-PLD would influence responsiveness to opioids and natural reinforcers. Given the precedent for sexual dimorphism in reward within both the eCB and opioid systems (Bradshaw et al., 2006; Collins et al., 2016; Gabel et al., 2022; Leishman et al., 2013), we asked how biological sex would affect results. To this end, we evaluated the performance of WT and NAPE-PLD KO mice in preclinical assays that address certain aspects of drug and natural reinforcement. First, we assessed WT and NAPE-PLD KO mice of both sexes in a locomotor sensitization paradigm to the opioid analgesic oxycodone. Next, we examined voluntary consumption of oral oxycodone in a continuous-access two bottle choice assay, tracking changes in body weight and temporal distribution of drinking activity in both genotypes and sexes. We also examined impact of oral oxycodone consumption on the number of ΔFosB-expressing cells in regions of both dorsal and ventral striata of WT and NAPE-PLD KO mice of both sexes. To investigate how NAPE-PLD influence natural avoidance and preference, we also tested male and female mice of both sexes in a two-bottle choice test for either quinine (a bitter taste) or sucrose. Our studies suggest that NAPE-PLD regulates responses to opioids in a sexually-dimorphic manner.

## 2. Methods

### 2.1 Animals

All procedures were approved by the Institutional Animal Care and Use Committee at Indiana University Bloomington. NAPE-PLD KO mice on a C57BL/6J background mice were bred as described previously (Leishman et al., 2016). NAPE-PLD KO mice in this study were generated either from a heterozygous cross (+/- with +/-) or as the F1 generation of a homozygous pair generated from the heterozygous cross. WT mice were supplemented with C57BL/6J mice purchased from Jackson Laboratories (Bar Harbor, ME, USA). Mice in each experiment were uniformly single- or group-housed (2-4 per cage, separated by sex) dictated by the experimental conditions. All mice had *ad libitum* access to water and food and were maintained on 12-hour light/dark cycle (lights on at 8:00 AM). All mice were between 12 and 16 weeks of age at time of testing.

### 2.2 Drugs and Chemicals

Oxycodone hydrochloride, sucrose, and quinine were purchased from Sigma-Aldrich (St. Louis, MO). For drug injection experiments, oxycodone was dissolved in 0.9% sterile saline and administered at an injection volume of 10 mL/kg (solution concentration 1.0 mg/mL for a 10 mg/kg injection). For oral consumption experiments, oxycodone, quinine, and sucrose were dissolved in sterile deionized water at the concentrations given in the text.

### 2.3 Behavioral Assays

Prior to all behavioral experiments, mice were handled by the experimenter in their colony room. All behavioral experiments were conducted during the animals’ light cycle between 10:00 AM and 3:00 PM.

#### 2.3.1 Locomotor Sensitization to Oxycodone

A 3-day protocol was used to examine acute hyperlocomotion as well as locomotor sensitization to oxycodone adapted from Severino and colleagues (Severino et al., 2020). Group-housed mice were marked on the tail with a Sharpie marker 1-2 days prior to behavioral testing for identification. Mice were brought into the room and habituated in their home cage on a table for 30-90 minutes prior to testing. For each session, mice were placed in activity meters (Superflex Nodes, Omnitech Electronics, Columbus, OH) for 30 minutes and locomotor activity was automatically recorded by photobeam interruptions interpreted by Fusion 6.5 software (Omnitech Electronics, Columbus, OH) starting the moment they entered the arena. The chambers were illuminated with tungsten bulbs at ∼80 lux, and a white noise generator provided a steady sound level of 62-63 dB in the arenas. In between testing each batch of animals, arenas were thoroughly cleaned with 70% ethanol before testing new animals in the same arena. The day prior to drug tests (Day 0), all mice were placed in the arena for 30 minutes to habituate them to the testing environment. On subsequent days (Days 1-3), mice received intraperitoneal (i.p.) injections of saline or oxycodone (10 mg/kg) and were placed into the arena immediately after injection. Mice were immediately returned to the colony room after testing.

#### 2.3.2 Home Cage Oral Oxycodone Two-Bottle Choice assay

We constructed open-source sipper devices which make use of inexpensive electronics, Arduino-based microcontrollers, and 3D printed parts to facilitate high-throughput assessment of drinking microstructure and fluid consumption (Godynyuk et al., 2019). These devices hold two 15 mL conical tubes (‘sippers’) with a drinking nozzle attached and quantify drinking behavior by measuring photobeam breaks at the nozzle of each sipper. To examine voluntary consumption of oral oxycodone, we adapted a continuous-access home cage two-bottle choice test as described previously (Slivicki et al., 2023). During the course of the experiment, mice were weighed daily (between 11AM and 3PM) and fluid volumes were evaluated manually using the side markings on the 15 mL conical vials (rounded to the nearest 0.25 mL). Daily fluid intake at each sipper tube was calculated as a function of body weight and presented as mg/kg (oxycodone) or ml/kg (water). Mice were randomly assigned to either an oxycodone group (oxycodone in treated sipper) or an opioid-naïve control (untreated water in the ‘treated’ sipper) and single housed a few days prior to the start of the experiment.

##### Habituation

Standard home-cage water bottles were replaced with sipper devices and mice were habituated to drinking from these devices with water in both tubes for three days prior to introduction of oxycodone.

##### Single Sipper Oxycodone Escalation Phase

During the first phase of this experiment, one sipper was removed from the device and mice were given escalating concentration of oxycodone for 6 days (Day 1: 0.1 mg/mL, Day 2-3: 0.3 mg/ml, Day 4-5: 0.5 mg/mL, Day 6: 1.0 mg/mL). The sipper’s position (right or left) was switched daily in an effort to teach mice to drink on both sides of the apparatus.

##### Two Bottle Choice Phase

On Day 7, the second bottle was returned to each mouse’s device, and mice were given the choice between their oxycodone-treated sipper (1.0 mg/ml) or an untreated bottle of water. During this phase, treated and untreated positions were fixed (i.e., stayed on either the left or right side) for the remaining 6 days to facilitate learning.

##### Seeking Test and Withdrawal

To test for drug-seeking behavior (Slivicki et al., 2023), sipper devices were removed from each mouse’s cage and replaced with standard issue water bottles containing untreated water after the 6^th^ day of the TBC phase. Then, the devices were returned to the cages after 24 hours of forced abstinence from oxycodone. This 1-hour test was performed with each sipper empty during the light cycle (at a time when drinking behavior is normally low). The number of beam breaks was quantified by the devices and serves as a proxy for number of attempts to drink from an empty sipper tube. Immediately after the seeking test (which was performed in the colony room), mice were brought to a testing room for analysis of locomotor activity in the activity meters described above. We recorded locomotor activity of control and oxycodone mice for a 15-minute period and additionally quantified the number of fecal boli left in the arena at the end of each session.

#### 2.3.3 Quinine and Sucrose Two Bottle Choice assay

To examine natural avoidance and preference, we examined a separate cohort of WT and NAPE-PLD KO mice of both sexes using a TBC paradigm similar to that performed by Reeves and colleagues (Reeves et al., 2021). During the entirety of this test, bottle sides were switched daily to eliminate side bias. Mice were given continuous home-cage access to the TBC device for the first 6 days of the experiment to habituate them to drinking (Days 0-5). During this phase, both bottles contained only water. Starting on Day 6, one bottle’s contents were replaced quinine (0.01 mM). The concentration of quinine was escalated every two days to allow one day of drinking on each side of the apparatus. After the quinine test (8 days, Day 6-13), mice were given a 6-day washout period in which both bottles once again contained only water (Day 14-19). Starting on Day 20, an equivalent procedure to the quinine test was performed with sucrose in concentrations of 0.25, 0.5, 1, and 2% (weight/volume), with 2 days at each concentration (Day 20-27). Consumption of quinine during each two-day period was averaged to give a single value for each concentration and compared across genotypes and sexes.

#### 2.3.4 Immunohistochemical staining of ΔFosB positive cells in the striatum

ΔFosB was quantified as previously described by our group (Iyer et al., 2021) by an experimenter (AB) blinded to all experimental conditions. Immediately after assessment of locomotor activity, used to measure possible spontaneous withdrawal, mice were deeply anesthetized with isoflurane, and transcardially perfused with 0.1% heparinized 0.1 M phosphate-buffered saline (PBS) followed by ice cold 4% paraformaldehyde (PFA). Brains were extracted, post-fixed for 24 hours in PFA, stored in PBS containing 0.1% sodium azide, and cryoprotected in 30% sucrose for three days prior to sectioning. Due to the variability in oxycodone consumption owing to the voluntary nature of the assay, brains were only sectioned from the 5-6 mice (i.e. 50% of subjects in each group) from each sex/genotype that consumed the most oxycodone or water based upon the median split analysis. Serial coronal floating sections (40 µM) containing the ventral and dorsal striatum were collected in 0.1 M PBS antifreeze solution (containing 50% sucrose and polyvinylpyrrolidone in ethylene glycol) guided by the mouse brain atlas of Paxinos and Franklin (Paxinos & Franklin, 2008) and stored at -20 °C until use. For staining, free-floating sections were immersed in PBS containing 0.3% hydrogen peroxide and non-specific binding was removed by incubation (1 h) with 5% goat serum diluted in PBS. Next, sections were incubated (4 °C) with a rabbit anti-ΔFosB antibody (1:10000, D3S8 R, Cell Signaling Technology) in 0.3% Triton PBS for 48 h. The tissue was incubated in the presence of biotinylated goat anti-rabbit IgG followed by Vectastain elite ABC reagent (1:600, #PK6101, Vector Laboratories, Burlingame, USA). ΔFosB immunoreactive cells were visualized with the avidin-biotin peroxidase method using diaminobenzidine as a chromogen. Sections were washed with double-distilled water, slide mounted, air-dried, dehydrated, and cover-slipped with Permount® (Fisher Scientific, Waltham, MA, USA).

#### 2.3.5 Quantification of Δ*FosB immunoreactive cells*

Images were captured from slide-mounted sections using a Leica (Wetzlar, Germany) DM6B microscope and a Leica DFC9000GT digital camera. The specificity of the immunostaining was verified by omission of the primary antibody from the immunostaining protocols as reported previously by our group (Iyer et al., 2021). ΔFosB-expressing nuclei were counted at 5x magnification bilaterally using a computer-assisted image analysis system (LAS AF 2D, Leica). A square field (200 x 200 μm) was superimposed upon the captured image to serve as a reference area and the number of cells were quantified by an investigator blinded to experimental conditions (i.e. genotype, sex, and drug treatment) of the mouse. The number of ΔFosB-expressing cells was quantified from three sections per mouse and averaged to generate a single value for each brain region for each animal (i.e. from a total of 41 mice) for use in statistical analyses.

### 2.6 Statistical Analysis

Behavioral data were analyzed with the following parametric and non-parametric statistical approaches. Differences in genotype, sex and consumption and their interactions were analyzed in behavioral and immunohistochemical studies using Two-way Analysis of Variance (ANOVA) followed by a Bonferroni post hoc test for multiple comparisons. All analyses were performed using GraphPad Prism version 10 (GraphPad Software, La Jolla, CA, USA). All data are presented as mean ± SEM unless otherwise stated. In all instances, p<0.05 was considered statistically significant. In figures, main effects of Two-way ANOVAs are presented under the following conventions: *p<0.05, **p<0.01, ***p<.001, ****p<.0001.

## 3. Results

### 3.1 **Figure 1**: Sex differences in the hyperlocomotive response to oxycodone present in WT mice are absent in NAPE-PLD KO mice

In response to the first injection of oxycodone (10 mg/kg, i.p.), female WT mice traveled a greater distance than male WT mice (**Fig 1A**) [Sex: F (1, 14) = 9.140, P=0.0091; Time: F (5, 70) = 1.852, P=0.1140; Sex x Time: F (5, 70) = 1.498, P=0.2017]. However, we did not observe sex differences in the acute hyperlocomotive response to oxycodone (10 mg/kg, i.p.) in NAPE-PLD KO mice (**Fig 1B**) [Sex: F (1, 16) = 0.03061, P=0.8633; Time: F (5, 80) = 20.37, P<0.0001; Sex x Time: F (5, 80) = 0.3462, P=0.8832]. Daily injections of oxycodone (10 mg/kg/day x 3 days, i.p.) increased locomotor activity across days in WT mice of both sexes, with females trending towards a greater locomotor response than males across sessions (**Fig 1C**) [Sex: F (1, 14) = 4.469, P=0.0529; Session: F (2, 28) = 56.23, P<0.0001; Sex x Session: F (2, 28) = 1.167, P=0.3258]. Daily injections of oxycodone (10 mg/kg/day x 3 days, i.p.) similarly increased distance traveled across sessions in NAPE-PLD KO mice, but no sex differences were observed in sensitization (**Fig 1D**) [Sex: F (1, 16) = 0.03889, P=0.8461; Session: F (2, 32) = 43.31, P<0.0001; Sex x Session: F (2, 32) = 1.143, P=0.3314]. When comparing genotypic sensitization responses within each sex across sessions, female NAPE-PLD KO mice treated with oxycodone exhibited a persistent reduction in distance travelled compared to oxycodone-treated female WT mice (**Fig 1E**) [Genotype: F (1, 16) = 13.41, P=0.0021; Session: F (2, 32) = 33.14, P<0.0001; Genotype x Session: F (2, 32) = 0.5284, P=0.5946]. In contrast, no reliable genotypic difference was observed in oxycodone-treated males under the same conditions (**Fig 1F**) [Genotype: F (1, 14) = 1.896, P=0.1902; Session: F (2, 28) = 76.51, P<0.0001; Genotype x Session: F (2, 28) = 0.6085, P=0.5512]. Both male (**Fig 1G**) and female (**Fig 1H**) saline-treated controls exhibited no genotypic difference in distance traveled across sessions [Males: Genotype: F (1, 14) = 0.1121, P=0.7428; Session: F (2, 28) = 1.002, P=0.3800; Genotype x Session: F (2, 28) = 1.638, P=0.2124. Females: Genotype: F (1, 14) = 3.666, P=0.0762; Session: F (2, 28) = 5.499, P=0.0097; Genotype x Session: F (2, 28) = 0.001652, P=0.9983].

**Figure 1:**
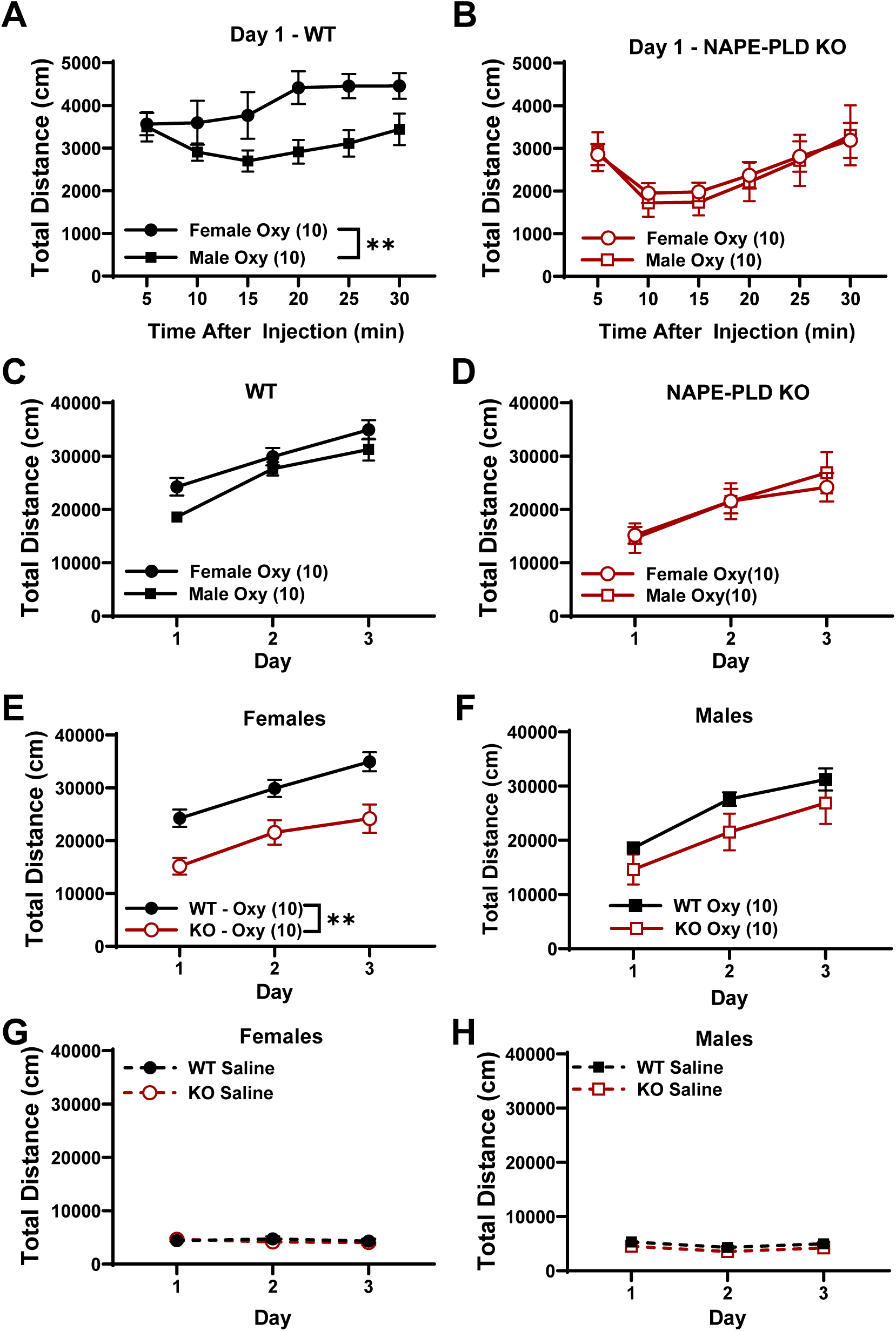
Sex differences in the hyperlocomotive response to oxycodone are observed in WT but not NAPE-PLD KO mice. In response to the first injection of oxycodone (10 mg/kg, i.p.) female WT mice traveled more distance than male WT mice **(A)**. No sex differences were observed in the acute hyperlocomotive response to oxycodone (10 mg/kg, i.p.) in NAPE-PLD KO mice **(B)**. Oxycodone (10 mg/kg/day x 3 days, i.p.) increased locomotor activity across sessions in WT mice of both sexes, with females trending towards a greater response than males across sessions **(C)**. NAPE-PLD KO mice increased their distance traveled across sessions but no sex differences were observed in the sensitization **(D)**. When comparing genotypic sensitization responses within each sex across sessions, female NAPE-PLD KO mice treated with oxycodone exhibited a persistent reduction in distance travelled compared oxycodone-treated female WT mice **(E)**. In contrast, no reliable genotypic difference was observed in oxycodone-treated males under the same conditions **(F)**. No genotypic difference in distance traveled was observed across sessions in male and female saline-treated controls **(G, H)**. Data are presented as the mean ± S.E.M. Bonferonni’s multiple comparisons: *p<0.05, **p<0.01. Sample size: N= 8-10 per sex per genotype per drug group.

### 3.2 Female NAPE-PLD KO mice consumed more oxycodone than both WT controls and male NAPE-PLD KO mice

All oxycodone-consuming mice escalated their oxycodone consumption as a function of concentrations given them during the first phase of the test. As reported by Slivicki and colleagues (Slivicki et al., 2023), we did not find reliable group differences between male and female WT mice in oxycodone consumption throughout the single or two bottle choice phases of the experiment (**Fig 2A**) [Sex: F (1, 22) = 1.429, P=0.2446; Day: F (11, 242) = 27.14, P<0.0001; Sex x Day: F (11, 242) = 0.4554, P=0.9287]. In contrast, we found that female NAPE-PLD KO mice consumed almost twice the quantity of oxycodone as male NAPE-PLD KO mice under the same conditions (**Fig 2B**) [Sex: F (1, 21) = 23.79, P<0.0001; Day: F (11, 231) = 32.56, P<0.0001; Sex x Day: F (11, 231) = 3.996, P<0.0001]. In within-sex comparisons, female NAPE-PLD KO mice consumed higher doses of oxycodone than female WT mice (**Fig 2C**) [Genotype: F (1, 21) = 9.744, P=0.0052; Day: F (11, 231) = 35.10, P<0.0001; Genotype x Day: F (11, 231) = 2.351, P=0.0092], while male mice did not differ in consumption by genotype (**Fig 2D**) [Genotype: F (1, 22) = 0.02633, P=0.8726; Day: F (11, 242) = 24.77, P<0.0001; Genotype x Day: F (11, 242) = 0.3774, P=0.9639]. Increased oxycodone consumption by female NAPE-PLD KO mice was mediated by a selective increase in consumption at the oxycodone-treated sipper (**Fig 2E**) [Genotype: F (1, 21) = 6.392, P=0.0195; Day: F (5, 105) = 2.857, P=0.0185; Genotype x Day: F (5, 105) = 0.8755, P=0.5003], while no non-specific increase in fluid consumption was observed at the untreated sipper containing just water (**Fig 2F**) [Genotype: F (1, 21) = 0.1910, P=0.6666; Day: F (5, 105) = 1.824, P=0.1143; Genotype x Day: F (5, 105) = 0.7042, P=0.6215]. Male WT and NAPE-PLD KO mice did not differ in consumption among males either at the oxycodone-treated sipper (**Fig 2G**) [Genotype: F (1, 22) = 0.1273, P=0.7247; Day: F (5, 110) = 0.3234, P=0.8979; Genotype x Day: F (5, 110) = 0.2666, P=0.9305] or at the untreated sipper (**Fig 2H**) [Genotype: F (1, 22) = 0.6542, P=0.4273; Day: F (5, 110) = 0.2411, P=0.9434; Genotype x Day: F (5, 110) = 0.6070, P=0.6946].

**Figure 2:**
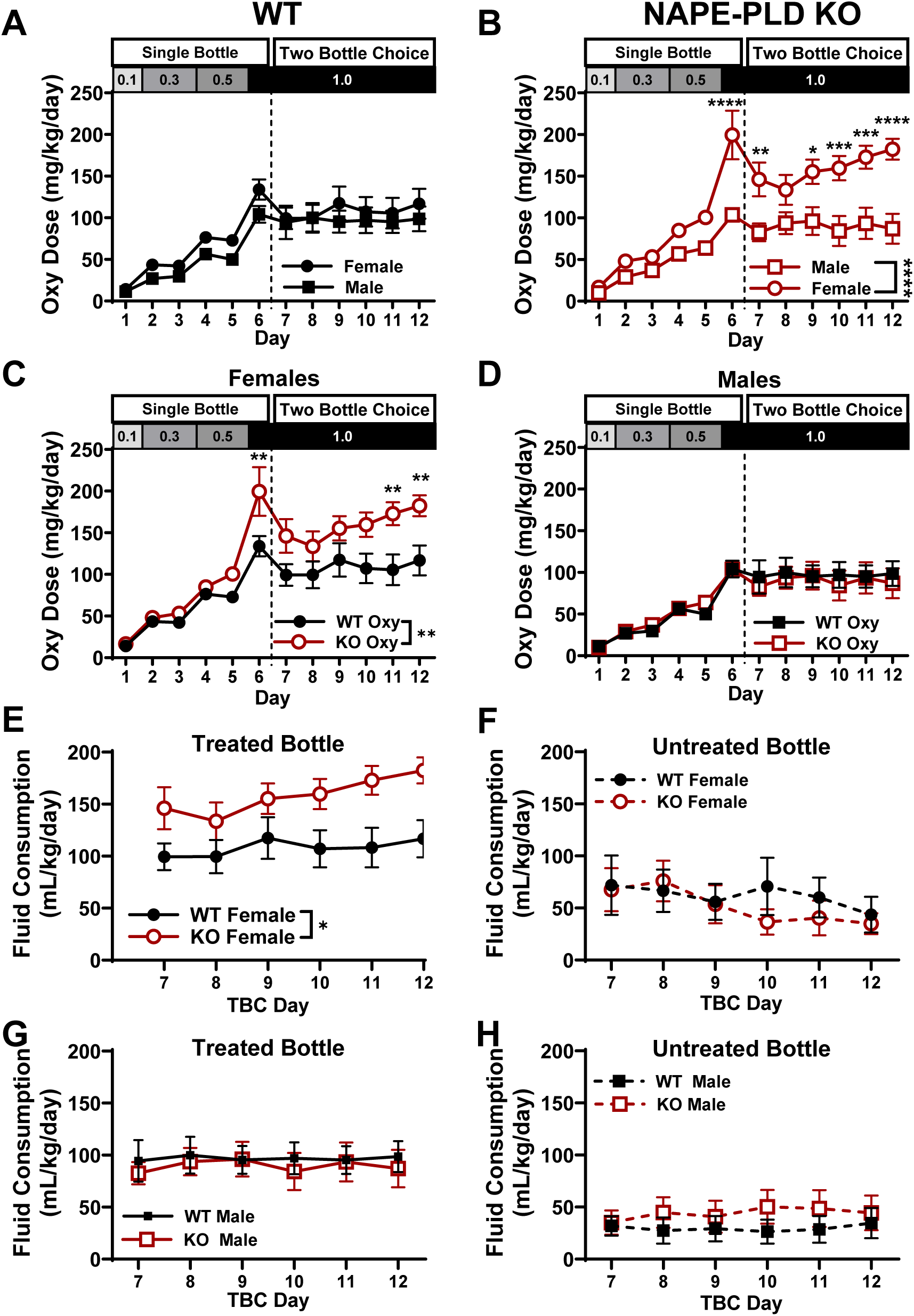
Female NAPE-PLD KO mice consumed more oxycodone than both WT controls and male NAPE-PLD KO mice. Oxycodone consumption increased as a function of the oxycodone concentrations presented during the first phase of the test (oxycodone concentration at each experimental timepoint at the top of **A-D**). Male and female WT mice did not differ in oxycodone consumption throughout the test **(A)**. Female NAPE-PLD KO mice consumed more oxycodone than male NAPE-PLD KO mice under the same conditions **(B)**. In within-sex comparisons, female NAPE-PLD KO mice consumed significantly higher doses of oxycodone than WT females **(C)**, while male mice did not differ in consumption by genotype **(D)**. In females, the genotypic difference in consumption was mediated by a selective increase in consumption at the oxycodone-treated sipper **(E)**, while no non-specific increase was observed at the untreated sipper containing just water **(F)**. We observed no reliable genotypic differences in consumption among males both at the oxycodone-treated sipper **(G)** as well as the untreated sipper **(H)**. Data are presented as the mean ± S.E.M. Bonferonni’s multiple comparisons: *p<0.05, **p<0.01, ***p<.001, ****p<.0001 for each comparison. Sample size: N=11-12 mice per sex per genotype.

### 3.3 Sex differences in weight loss from oxycodone consumption are reduced in NAPE-PLD KO mice

An experimental timeline referencing the experimental phases and oxycodone concentrations available for consumption on each day are shown in **Fig 3A**. Female WT mice that consumed only water from the sipper device gained more bodyweight (approximately 5%) compared to those that consumed oxycodone **(Fig. 3B)** [Drug: F (1, 20) = 6.937, P=0.0159; Day: F (11, 220) = 8.023, P<0.0001; Drug x Day: F (11, 220) = 1.750, P=0.0643]. In contrast, compared to water-drinking controls (Control Male), WT males that consumed oxycodone reliably lost bodyweight (<10%) during the course of the experiment **(Fig 3C)** [Drug: F (1, 20) = 71.12, P<0.0001; Day: F (11, 220) = 10.63, P<0.0001; Drug x Day: F (11, 220) = 23.17, P<0.0001]. A comparison between sexes revealed that male WT mice that consumed oxycodone lost weight over the course of the study whereas female WT mice were largely unaffected **(Fig 3D)** [Sex: F (1, 22) = 56.64, P<0.0001; Day: F (11, 242) = 8.958, P<0.0001; Sex x Day: F (11, 242) = 18.48, P<0.0001]. In contrast, oxycodone-drinking NAPE-PLD KO mice of both sexes lost weight, which was more prominent among males than females **(Fig 3E)** [Sex: F (1, 21) = 6.303, P=0.0203; Day: F (11, 231) = 24.76, P<0.0001; Sex x Day: F (11, 231) = 3.443, P=0.0002]. Female NAPE-PLD KO mice lost more bodyweight across time than female WT mice **(Fig 3F)** [Genotype: F (1, 21) = 29.89, P<0.0001; Day: F (11, 231) = 5.772, P<0.0001; Genotype x Day: F (11, 231) = 5.722, P<0.0001]. Male NAPE-PLD KO mice consuming oxycodone lost more weight loss compared to male WT mice consuming oxycodone. However, the genotypic difference was less in males than females (**Fig 3G**) [Genotype: F (1, 22) = 5.820, P=0.0246; Day: F (11, 242) = 49.16, P<0.0001; Genotype x Day: F (11, 242) = 1.786, P=0.0570].

**Figure 3:**
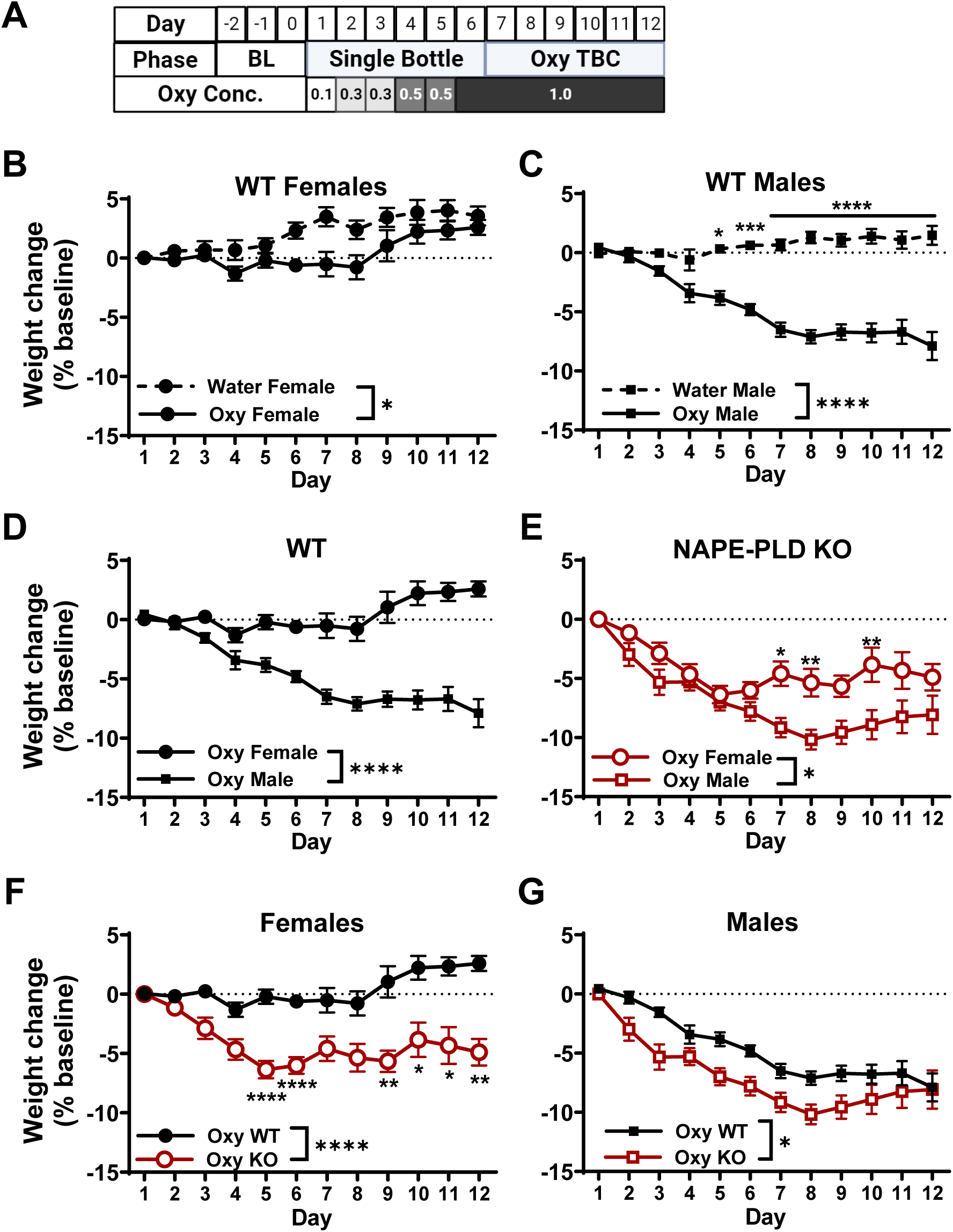
WT mice exhibit sex differences in weight loss following oxycodone consumption that are reduced in NAPE-PLD KO mice. An experimental timeline shows experimental phases and oxycodone concentrations available to mice on each day in a voluntary oral consumption paradigm **(A)**. Female WT mice that consumed only water (Water Female) gained weight, whereas weight gain was slightly delayed in mice that consumed oxycodone **(B)**. Compared to water-drinking controls (Water Male) oxycodone-drinking WT males lost body weight during the course of the experiment **(C)**. Only male WT mice exhibited reductions in body weight, whereas females were largely unaffected **(D)**. Oxycodone-drinking NAPE-PLD KO mice of both sexes exhibited weight loss which was less prominent in females **(E)**. Genotypic comparison among females reveals a weight drop in NAPE-PLD KO mice **(F)**. Compared to WT males, male NAPE-PLD KO mice exhibited a modest amplification of weight loss, though to a lesser extent than the genotypic difference seen in females **(G)**. Data are presented as the mean ± S.E.M. Bonferonni’s multiple comparisons: *p<0.05, **p<0.01, ***p<.001, ****p<.0001 for each comparison. Sample size: N=10-12 mice per sex per genotype per treatment group.

### 3.4 Drug-seeking behavior and signs of spontaneous withdrawal after forced abstinence from oxycodone were present in both genotypes

After the 6-day two bottle choice phase of the experiments, drinking devices were removed from the home cage and replaced with standard water bottles (see **Fig 4A** for schematic). Then, 24 hours later, mice were tested for oxycodone-seeking behavior by returning the device to the cage with empty bottles for 1 hour (with the alternative water bottle also available at this time). After 24 hours of forced abstinence, male, but not female, WT mice that previously consumed oxycodone trended to spend more time drinking from the empty sipper compared to water-drinking controls (**Fig 4B**) [Drug: F (1, 38) = 2.446, P=0.1261; Sex: F (1, 38) = 1.298, P=0.2616; Drug x Sex: F (1, 38) = 4.009, P=0.0524]. After 24 hours of forced abstinence, oxycodone-consuming NAPE-PLD KO mice of both sexes attempted to drink from an empty sipper more than water-drinking controls (**Fig 4C**) [Drug: F (1, 40) = 8.069, P=0.0070; Sex: F (1, 40) = 0.9608, P=0.3329; Drug x Sex: F (1, 40) = 0.4311, P=0.5152]. When evaluated in activity meters after the seeking test, WT mice with a history of oxycodone consumption traveled less distance than WT mice, and female WT mice traveled farther than males regardless of drug condition (**Fig 4D**) [Drug: F (1, 39) = 11.80, P=0.0014; Sex: F (1, 39) = 6.130, P=0.0177; Drug x Sex: F (1, 39) = 1.640, P=0.2079]. NAPE-PLD KO mice with a history of oxycodone consumption traveled less distance than their water-drinking counterparts, irrespective of sex (**Fig 4E**) [Drug: F (1, 40) = 5.070, P=0.0299; Sex: F (1, 40) = 1.116, P=0.2971; Drug x Sex: F (1, 40) = 0.07997, P=0.7788]. Prior oxycodone consumption increased the amount of time that WT (**Fig 4F**) and NAPE-PLD KO (**Fig 4G**) spent resting during the test [WT Fig 4F: Drug: F (1, 39) = 34.20, P<0.0001; Sex: F (1, 39) = 0.5926, P=0.4461; Drug x Sex: F (1, 39) = 0.005499, P=0.9413. NAPE-PLD KO Fig 4G: Drug: F (1, 40) = 0.002297, P=0.9620; Sex: F (1, 40) = 18.66, P=0.0001; Drug x Sex: F (1, 40) = 0.06786, P=0.7958]. Additionally, previous oxycodone consumption increased defecation in WT mice (**Fig 4H**) [Drug: F (1, 39) = 34.00, P<0.0001; Sex: F (1, 39) = 0.2029, P=0.6549; Drug x Sex: F (1, 39) = 0.1066, P=0.7458]. Oxycodone consumption similarly increased defecation in NAPE-PLD KO mice, while male NAPE-PLD KO mice defecated more than female KO mice overall (**Fig 4I**) [Drug: F (1, 40) = 9.920, P=0.0031; Sex: F (1, 40) = 4.724, P=0.0357; Drug x Sex: F (1, 40) = 1.339, P=0.2542].

**Figure 4:**
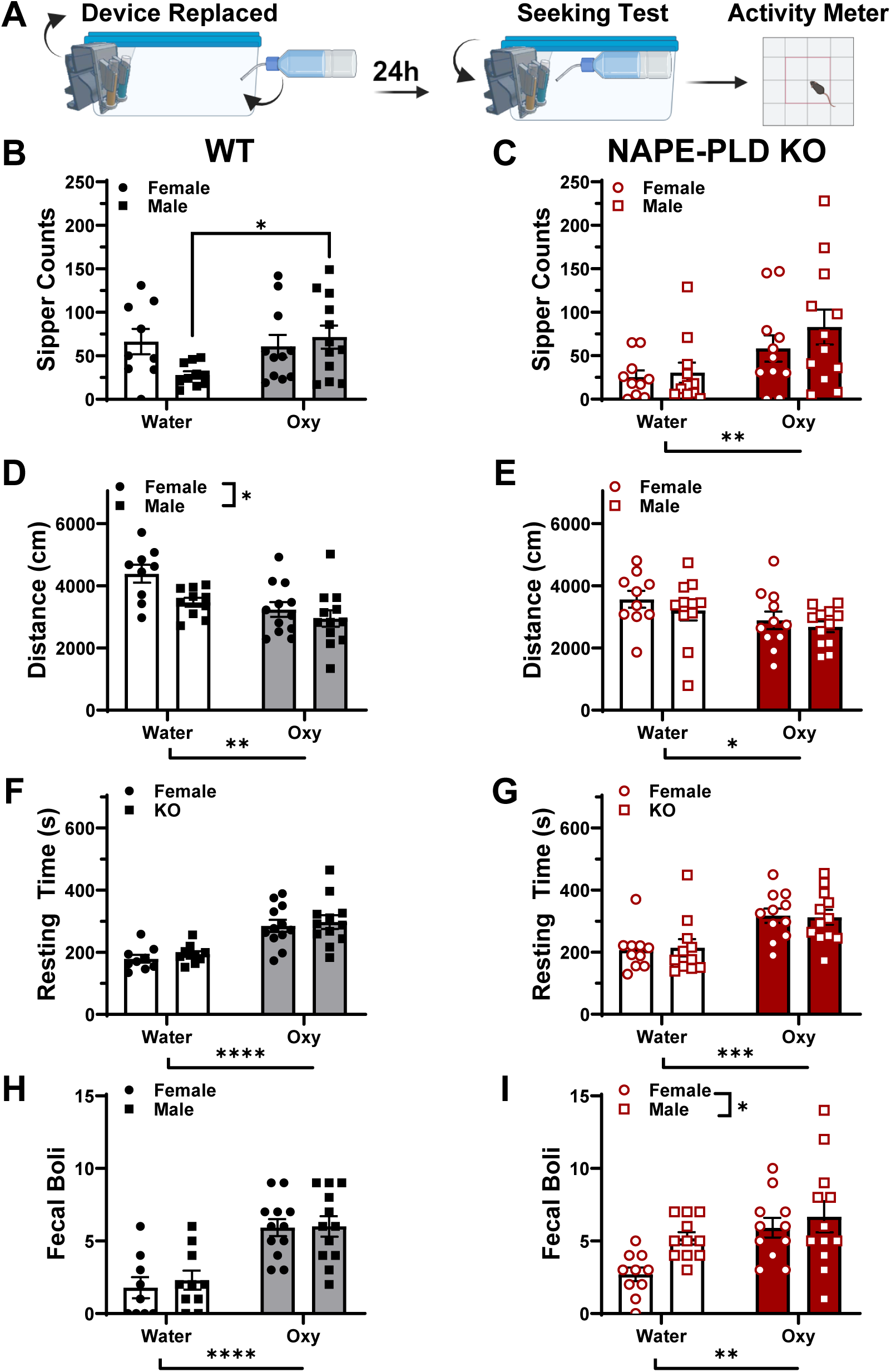
Drug-seeking behavior and signs of spontaneous withdrawal after forced abstinence from oxycodone were present in both genotypes. After 24 hours of forced abstinence, male but not female WT that had previously consumed oxycodone attempted to drink from an empty sipper more than water-drinking controls **(B).** NAPE-PLD KO mice of both sexes that previously consumed oxycodone attempted to drink from an empty sipper more than water-drinking controls **(C)**. When evaluated in activity meters after the seeking test, WT **(D)** and NAPE-PLD KO **(E)** oxycodone-drinking mice traveled less distance than water-drinking controls. WT females traveled more distance than males **(D)**. Prior oxycodone consumption increased the amount of time that WT **(F)** and NAPE-PLD KO **(G)** spent resting during the test. Additionally, previous oxycodone consumption increased defecation in WT **(H)** and NAPE-PLD KO **(I)** mice, while male NAPE-PLD KO mice defecated more than female KO mice overall **(I)**. Data are presented as the mean ± S.E.M. Bonferonni’s multiple comparisons: *p<0.05 for each comparison. Sample size: N=10-12 mice per sex per genotype per treatment group.

### 3.5 Expression of the transcription factor **Δ**FosB was selectively increased in the dorsolateral striatum of WT females that consumed oxycodone

After assessment of locomotor activity during forced abstinence in the seeking test, mice with a history of oxycodone or water consumption were perfused and brains removed for immunohistochemical analysis of ΔFosB. Representative photomicrographs showing immunolabeling for ΔFosB in coronal sections containing the dorsal and ventral striatum (see **Supplemental Fig 1** for depiction of regions quantified) are shown for WT (**Fig 5A-D**) and NAPE-PLD KO (**Fig 5E-H**) mice consuming either water (**Fig 5A, C, E, G**) or oxycodone (**Fig 5B, D, F, H**). Oxycodone consumption increased the number of ΔFosB positive cells in the dorsolateral striatum of female, but not male, WT mice (**Fig 6A**). Additionally, WT females exhibited more ΔFosB positive cells in the dorsolateral striatum than that observed in the same region of WT males (**Fig 6A**) [Drug: F (1, 16) = 6.064, P=0.0255; Sex: F (1, 16) = 4.691, P=0.0458; Drug x Sex: F (1, 16) = 9.633, P=0.0068]. Despite the elevated consumption of oxycodone observed in female NAPE-PLD KO mice, oxycodone consumption did not exhibit an increase in the number of ΔFosB positive cells in the dorsolateral striatum of NAPE-PLD KO mice of either sex (**Fig 6B**) [Drug: F (1, 17) = 1.156, P=0.2972; Sex: F (1, 17) = 1.142, P=0.3001; Drug x Sex: F (1, 17) = 0.02849, P=0.8680]. No differences were observed in the number of ΔFosB positive cells as a function of sex or drug group in the dorsomedial striatum of WT (**Fig 6C**) or NAPE-PLD KO mice (**Fig 6D**) [WT Fig 6C: Drug: F (1, 16) = 1.283, P=0.2741; Sex: F (1, 16) = 3.029, P=0.1010; Drug x Sex: F (1, 16) = 3.224, P=0.0915. NAPE-PLD KO Fig 6D: Drug: F (1, 17) = 1.832, P=0.1936; Sex: F (1, 17) = 1.005e-005, P=0.9975; Drug x Sex: F (1, 17) = 0.1329, P=0.7199]. In the nucleus accumbens core (**Fig 6E** and **6F**) and shell (**6G** and **6H**), we similarly observed no differences in quantity of ΔFosB positive cells as a function of sex or drug in WT (**Fig 6E** and **6G**) or NAPE-PLD KO mice (**Fig 6F** and **6H**) [WT Core Fig 6E: Drug: F (1, 16) = 0.1297, P=0.7234; Sex: F (1, 16) = 0.04845, P=0.8286; Drug x Sex: F (1, 16) = 1.211, P=0.2874. NAPE-PLD KO Core Fig 6F: Drug: F (1, 17) = 0.6826, P=0.4201; Sex: F (1, 17) = 0.1320, P=0.7209; Drug x Sex: F (1, 17) = 3.346, P=0.0850. WT shell Fig 6G: Drug: F (1, 16) = 0.6884, P=0.4189; Sex: F (1, 16) = 0.7984, P=0.3848; Drug x Sex: F (1, 16) = 0.8763, P=0.3631.NAPE-PLD KO shell Fig 6H: Drug: F (1, 17) = 0.07624, P=0.7858; Sex: F (1, 17) = 0.01752, P=0.8962; Drug x Sex: F (1, 17) = 1.823, P=0.1946].

**Figure 5.**
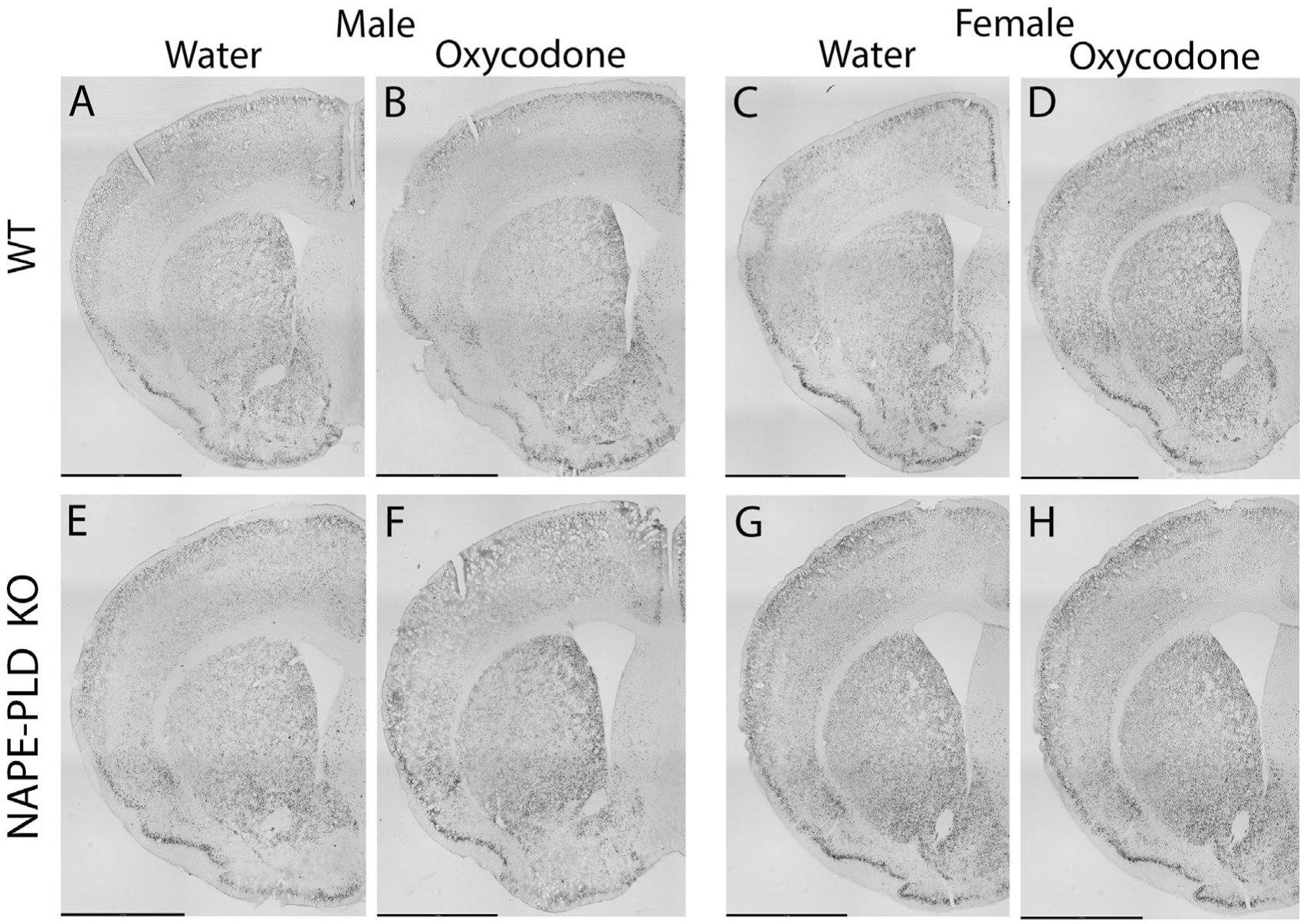
Representative images showing mmunohistochemical staining for ΔFosB i in mouse coronal sections. Photomicrographs show coronal sections at the level of the striatum that were stained for quantification of ΔFosB. Representative images are shown for male **(A, B)** and female **(C, D)** WT mice that consumed only water **(A, C)** or oxycodone **(B, D)**, and male **(E, F)** and female **(G, H)** NAPE-PLD KO mice that consumed water (**E, G)** or oxycodone **(F, H)**. Scale bar represents1.7 mm.

**Figure 6.**
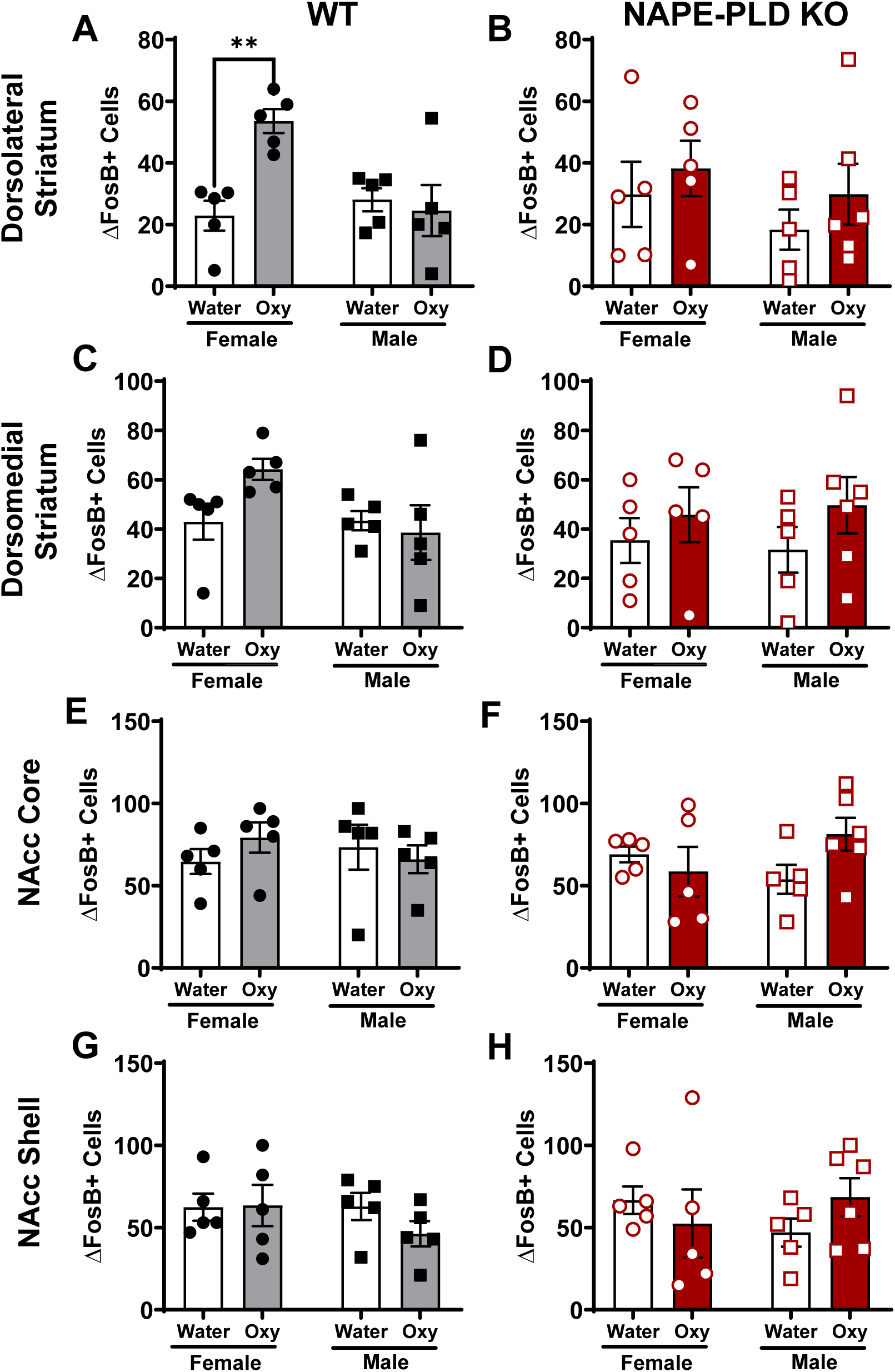
Voluntary consumption and forced abstinence from oxycodone elevated levels of ΔFosB selectively in the dorsolateral striatum of WT female mice. Oxycodone consumption increased the number of ΔFosB-expressing cells in the dorsolateral striatum of high consuming female but not male WT mice (**A**). Oxycodone consumption did not affect the number of ΔFosB expressing cells among NAPE-PLD KO mice of either sex within this brain region (**B**). No differences in the number of ΔFosB-expressing cells were observed as a function of drug or sex in any other brain region surveyed, including the dorsomedial striatum (**C**, **D**), the nucleus accumbens (NAcc) core (**E**, **F**), and the NAcc Shell (**G**, **H**) in either WT (**C, E, G**) or NAPE-PLD KO (**D, F, G**) mice. Three sections per mouse were quantified for each brain region and averaged into a single mean for each mouse. Data are presented as the mean ± S.E.M. Two-way ANOVA followed by Bonferonni’s multiple comparisons: **p<0.01 for each comparison. Sample size: N=5-6 mice per group.

### 3.6 Home cage consumption of sucrose, but not quinine, was amplified in NAPE-PLD KO mice while sex differences in sucrose consumption were preserved

Consumption of quinine in a two-bottle choice test did not differ in either females (**Fig 7A**) or males (**Fig 7B**) as a function of genotype at any concentration [Males **Fig 7A**: Concentration: F (1.298, 18.17) = 7.100, P=0.0109; Genotype: F (1, 14) = 0.5408, P=0.4742; Concentration x Genotype: F (3, 42) = 0.2995, P=0.8256. Female **Fig 7B**: Concentration: F (1.056, 14.78) = 4.246, P=0.0557; Genotype: F (1, 14) = 0.5486, P=0.4711; Concentration x Genotype: F (3, 42) = 1.073, P=0.3710]. However, both female (**Fig 7C**) and male (**Fig 7D**) NAPE-PLD KO mice consumed more sucrose than WT controls at the highest concentration of sucrose [**Female Fig 7C**: Concentration: F (3, 42) = 4.052, P=0.0129; Genotype: F (1, 14) = 3.493, P=0.0827; Concentration x Genotype: F (1.109, 15.52) = 138.9, P<0.0001. **Male Fig 7D**: Concentration: F (1.202, 16.83) = 71.81, P<0.0001; Genotype: F (1, 14) = 3.565, P=0.0799; Concentration x Genotype: F (3, 42) = 5.614, P=0.0025]. Female WT mice consumed more sucrose overall than male WT mice, particularly at the highest concentrations (1% and 2%) of sucrose (**Fig 7E**) [Concentration: F (1.236, 17.31) = 121.5, P<0.0001; Sex: F (1, 14) = 17.82, P=0.0009; Concentration x Sex: F (3, 42) = 19.50, P<0.0001]. The sex difference in sucrose consumption was also observed in NAPE-PLD KO mice (**Fig 7F**) [Concentration: F (1.087, 15.22) = 105.3, P<0.0001; Sex: F (1, 14) = 19.82, P=0.0005; Concentration x Sex: F (3, 42) = 9.581, P<0.0001]. By contrast, a cross-sectional comparison of consumption at the highest concentration of quinine revealed no sex or genotypic differences in quinine consumption (**Fig 6G**) [Sex: F (1, 28) = 0.7048, P=0.4083; Genotype: F (1, 28) = 1.051, P=0.3140; Sex x Genotype: F (1, 28) = 0.05394, P=0.8180]. In contrast, a similar analysis of sucrose consumption at the highest concentration tested reveals that NAPE-PLD KO mice consumed more sucrose that WT mice, and females consumed more than males irrespective of genotype (**Fig 6H**) [Sex: F (1, 28) = 29.78, P<0.0001; Genotype: F (1, 28) = 9.242, P=0.0051; Sex x Genotype: F (1, 28) = 0.1606, P=0.6916].

**Figure 7.**
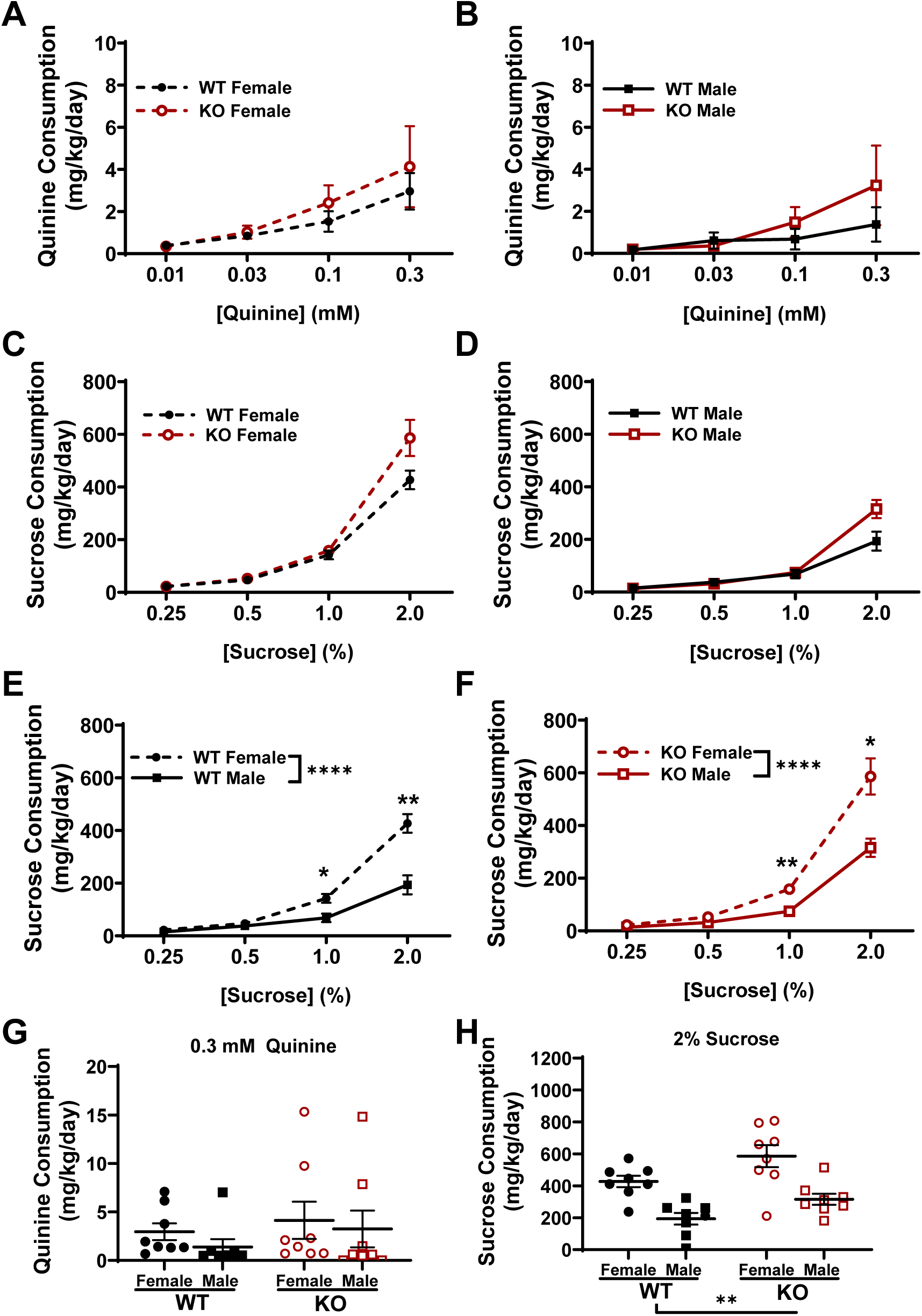
Home cage consumption of sucrose, but not quinine, was increased in NAPE-PLD KO mice compared to WT mice whereas sex differences in sucrose consumption were preserved in both genotypes. In a two-bottle choice test, consumption of quinine did not differ as a function of genotype at any concentration in either female **(A)** or male **(B)** mice. Both female **(C)** and male **(D)** NAPE-PLD KO mice consumed more sucrose than WT controls with increasing concentrations of sucrose [Female Interaction: P<0.0001; Male Interaction: P=0.0025]. Females consumed more sucrose than males among both WT **(E)** and NAPE-PLD KO mice **(F)**. No sex or genotypic differences in quinine consumption were observed at the highest concentration of quinine based upon a cross-sectional sex by genotype comparison **(G)**. In contrast, females of both genotypes consumed more sucrose than males at the 2% concentration **(H)**. Data are presented as the mean ± S.E.M. Bonferroni’s multiple comparisons: *p<0.05, **p<0.01, ***p<.001, for each comparison. Sample size: N=8 mice per sex per genotype.

## 4. Discussion

Here we show that genetic deletion of NAPE-PLD influenced responsiveness to oxycodone in a sexually dimorphic and context-dependent manner. We observed sexual dimorphism in both oxycodone-induced hyperlocomotion and weight loss of WT mice, as others have previously observed (Collins et al., 2016). However, sex differences in the hyperlocomotor response to oxycodone were virtually absent in NAPE-PLD KO mice. Conversely, while sex differences were absent in WT mice in a voluntary oral oxycodone consumption paradigm, female NAPE-PLD KO mice consumed twice the amount of oxycodone as male KO mice. Finally, sex differences in consumption of sucrose (i.e., heightened consumption among females) were observed in both WT and NAPE-PLD KO mice.

WT mice consuming oxycodone lost more bodyweight compared to control mice consuming water and this effect was exacerbated in NAPE-PLD KO mice. In WT mice, voluntary oxycodone consumption produced weight loss that was preferentially observed in males (despite equivalent group consumption between sexes), while oxycodone-consuming NAPE-PLD KO mice of both sexes that consumed oxycodone lost weight. This observation could be due to differential sensitivity in males and females in the dose required to disrupt the homeostasis of body weight. As female NAPE-PLD KO mice consumed almost twice the dose of oxycodone as female WT mice, they may have reached that threshold. This finding potentially aligns with previous findings that female mice are less sensitive than male mice to weight loss in response to chronic, high doses of morphine (Li et al., 2022). We did not observe any reliable changes in bodyweight in response to 3 daily injections of oxycodone at 10 mg/kg i.p. in our locomotor sensitization paradigm (data not shown), alluding to the dose-dependent nature of this effect.

We previously reported that voluntary consumption of oxycodone in a limited access two-bottle choice test produced physical dependence, as measured by quantification of naloxone-precipitated opioid withdrawal jumps, which correlated with the number of ΔFosB expressing cells in the nucleus accumbens (NAc) shell (Iyer et al., 2021). ΔFosB is a transcription factor whose accumulation plays an important role in altering brain plasticity in response to chronic drug exposure (Nestler, 2008). In the spontaneous withdrawal paradigm employed here, the number of ΔFosB expressing cells in the ventral striatum were not affected by oral oxycodone consumption. This discrepancy could be in part due to differences in experimental parameters, including the dose of oxycodone consumed in the present continuous access paradigm (vs. previous limited access paradigm), assessment of spontaneous withdrawal (vs. naloxone-precipitated opioid withdrawal), the timepoint of perfusion and dissection, and small sample size. We did, however, observe increases in the number of ΔFosB-expressing cells in the dorsolateral striatum of female, but not male, WT mice that consumed oxycodone, while no differences were observed in female relative to male NAPE-PLD KO mice despite their elevated consumption. This observation is of interest because it suggests that the reduced behavioral sensitivity to oxycodone observed in female NAPE-PLD KO mice extended to the immunohistochemical level as well. While exogenously administered AEA elevated levels of ΔFosB in one study (Salaya-Velazquez et al., 2020), little is presently known about how endogenous lipid signaling via NAEs affects ΔFosB accumulation.

Across paradigms, NAPE-PLD deletion altered the presentation of the sexually dimorphic response to oxycodone. Several factors are known to mediate sex differences in opioid responsivity, including drug metabolism, sex hormones, and expression/activity of opioid receptors. Previous studies suggest that estrous cycle influences analgesia and i.v. self-administration of oxycodone, with females in diestrus exhibiting enhanced analgesia and self-administration of oxycodone (Arguelles et al., 2021; Hinds et al., 2023). Interestingly, orally-administered oxycodone levels were higher in the brains of females in diestrus compared to males and females in estrus, potentially due to diminished brain activity of the cytochrome P450 (CYP) enzyme CYP2D during diestrus (Arguelles et al., 2021). While NAPE-PLD is known to play a role in lipid metabolism (Geurts et al., 2015; Lefort et al., 2020), it is unclear if NAEs affect the activity or expression of cytochrome P450 (CYP) enzymes that metabolize oxycodone. Nonetheless, NAPE-PLD is expressed in the murine and human female reproductive tract, with its quantities/products fluctuating depending on stage of menstrual cycle or pregnancy (Scotchie et al., 2015; Wang et al., 2008). Brain levels of anandamide and congeners are also known to fluctuate depending on stage of estrous cycle (Bradshaw et al., 2006). The primary objective of the present study was to characterize behavioral sensitivity to oxycodone in male and female WT/NAPE-PLD KO mice. Determining if alterations in the sexually dimorphic drug response in NAPE-PLD KO mice are mediated by NAPE-PLD’s influence on levels of sex hormones/estrous cycle could be an excellent mechanistic follow up for the present study, which would require experimental manipulation of hormones and tracking of estrous cycle.

In the present study, NAPE-PLD KO mice consumed higher quantities of sucrose than WT controls of both sexes in a continuous access two bottle choice paradigm. This is somewhat at odds with recently published findings from a separate NAPE-PLD KO line, where NAPE-PLD KO mice exhibited less sucrose preference than WT mice (Chen et al., 2023). Genetic differences in the two KO lines and/or distinct behavioral contexts may underlie this discrepancy. For example, due to the genetic approach used to generate the NAPE-PLD KO line used Chen et al., 2023 (known as the Cravatt line), these knockouts exhibit reductions in some NAEs but not AEA (Liu et al., 2008; Mock et al., 2020). However, the Luquet line used here, which removed the catalytic site in exon 3, exhibits reductions in AEA and all measured NAEs in all brain regions compared to WT mice (Leishman et al., 2016). Additionally, the sucrose preference test utilized in Chen et al 2023 involved 24h water deprivation, which potentially could have caused an acute stress response (Bekkevold et al., 2013). This is important because we have characterized dysregulation of the HPA axis in the Luquet line (Woodward et al., 2024), and the sucrose preference test in this assay would presumably be less stressful as a continuous access paradigm. Regardless of the genetic or procedural differences in our findings, future studies should directly compare the two NAPE-PLD KO lines to examine how a reduction in non-anandamide NAEs (Cravatt line) would influence behavioral outcomes versus reduction of AEA as well as all NAEs (Luquet line). Conversely, another group recently found that conditional deletion of NAPE-PLD in neurons or locally in the ventral tegmental area enhanced consumption of sucralose and sucrose and in general enhances dopamine release in response to both food and non-food related reward (Castel et al., 2024) similar to what was observed in our study. More work is necessary to determine whether phenotype of NAPE-PLD KO mouse revealed here would be recapitulated with region-specific deletion of NAPE-PLD in adulthood in future studies elucidating possible contributions of central (e.g. VTA, nucleus accumbens) and peripheral NAPE-PLD. The differences and similarities found in these studies speak to the complexity of lipid metabolism, the effects of different genetic approaches, and the tissue-dependent and context-dependent manner through which NAPE-PLD affects sensitivity to drugs and natural rewards.

One caveat in interpreting the results of the present study is that the WT mice used in this study were not littermates of NAPE-PLD KO mice. Because of the large number of mice required to fully evaluate sex differences and drug effects in each genotype, we opted for this strategy for our initial investigations. This approach minimized the number of mice that would otherwise be unnecessarily bred (>100 heterozygous littermates would be generated, and not used for data collection) and, consequently, killed. Given this caveat, we provide within-genotype characterizations of sex differences and drug effects with saline/water controls, which provides important new information even with this stated limitation.

Together, these findings lay groundwork for future studies of surrounding the interactions of NAPE-PLD activity, drug reinforcement, and sex differences. Future studies employing conditional and/or inducible knockout strategies will also shed light into which cell types and developmental effects contribute to the observed results in our study. Careful evaluation and manipulation of sex hormones in future work may offer insights into mechanisms of alterations in sexually dimorphic drug response that we observed.

## Acknowledgements

Graphics for experimental schematics were created using biorender.com.

## Authors’ contributions

TJW, ES, AB, AP, JH, and HK – investigation and formal analysis. TJW and AGH – conceptualization. TJW, AP, AB, EF – Visualization. AGH –supervision and project administration. TJW and AGH –Writing-original draft. KM, ES, AB, and SL– Writing-review and editing. AGH, SL and KM-resources.

---

**Supplementary Figure 1.**
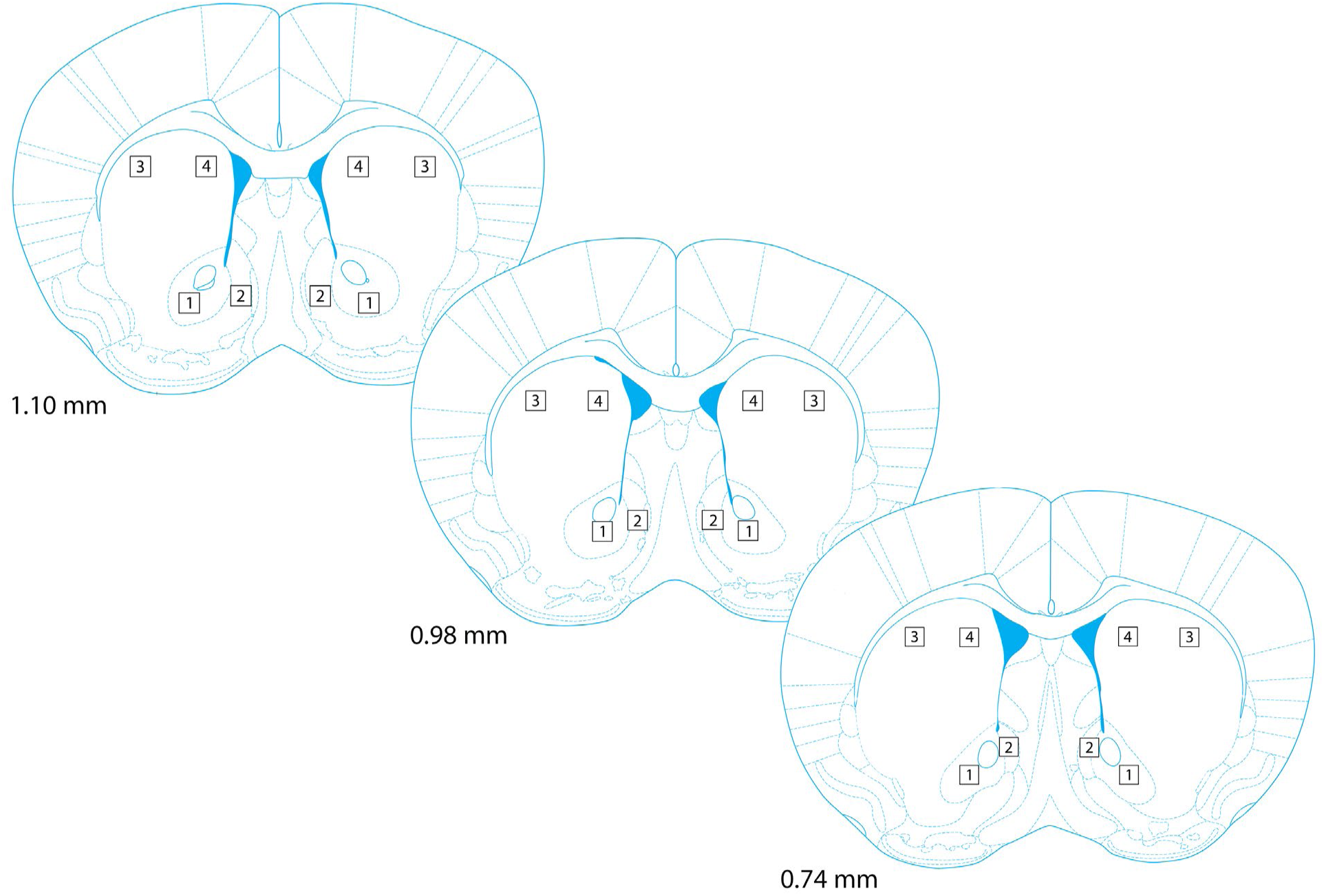
Schematic showing levels of coronal sections and brain regions quantified for ΔFosB expression levels. A 200 µm x 200 µm field was used for quantification of ΔFosB immunopositive cells in the nucleus accumbens core (Box 1), nucleus accumbens shell (Box 2), dorsolateral striatum (Box 3), and dorsomedial striatum (Box 4) at different coronal levels. Schematics show distance (in mm) of coronal plane of section relative to bregma (Paxinos and Franklin (2008). Quantification was based upon 3 sections per mouse.

